## Supplementary materials for "Intrinsically disordered pathogen effector alters the STAT1 dimer to prevent recruitment of co-transcriptional activators CBP/p300"

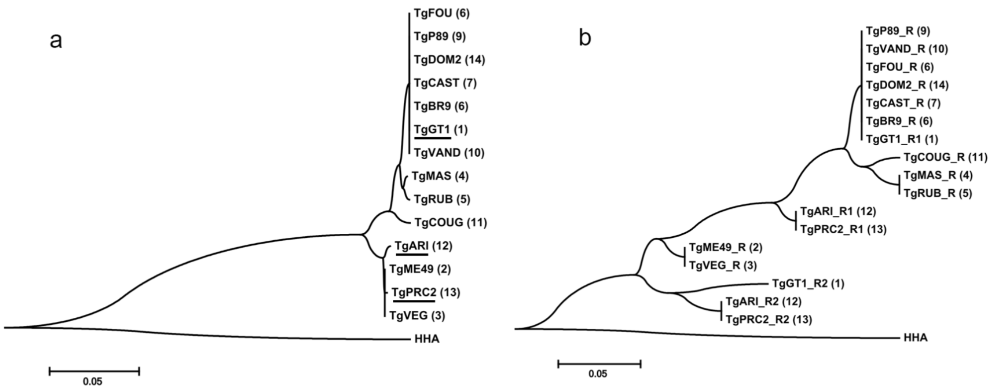


**Fig. S1**

**Phylogenetic analysis ofTgIST from different lineages of *T.gondii*.** (a) Neighbor-joining tree of full length TgIST from *T. gondii* strains (TG) or *Hammondia hammondi* (HHA). Strain types are indicated in the parentheses. Strains with a duplciated repeat region are underlined. (b) Neighbor-joining tree of individual repeat regions within TgIST from *T. gondii* strains (TG) or *Hammondia hammondi* (HHA). Protein sequences of TgIST were retrived from ToxoDB (https://www.toxodb.org/) and aligned using Muscle for mutiple sequence alignments ( https://www.ebi.ac.uk/Tools/msa/muscle/). Phylogenetic trees were viualized by MEGA 7.0 (https://www.megasoftware.net/).

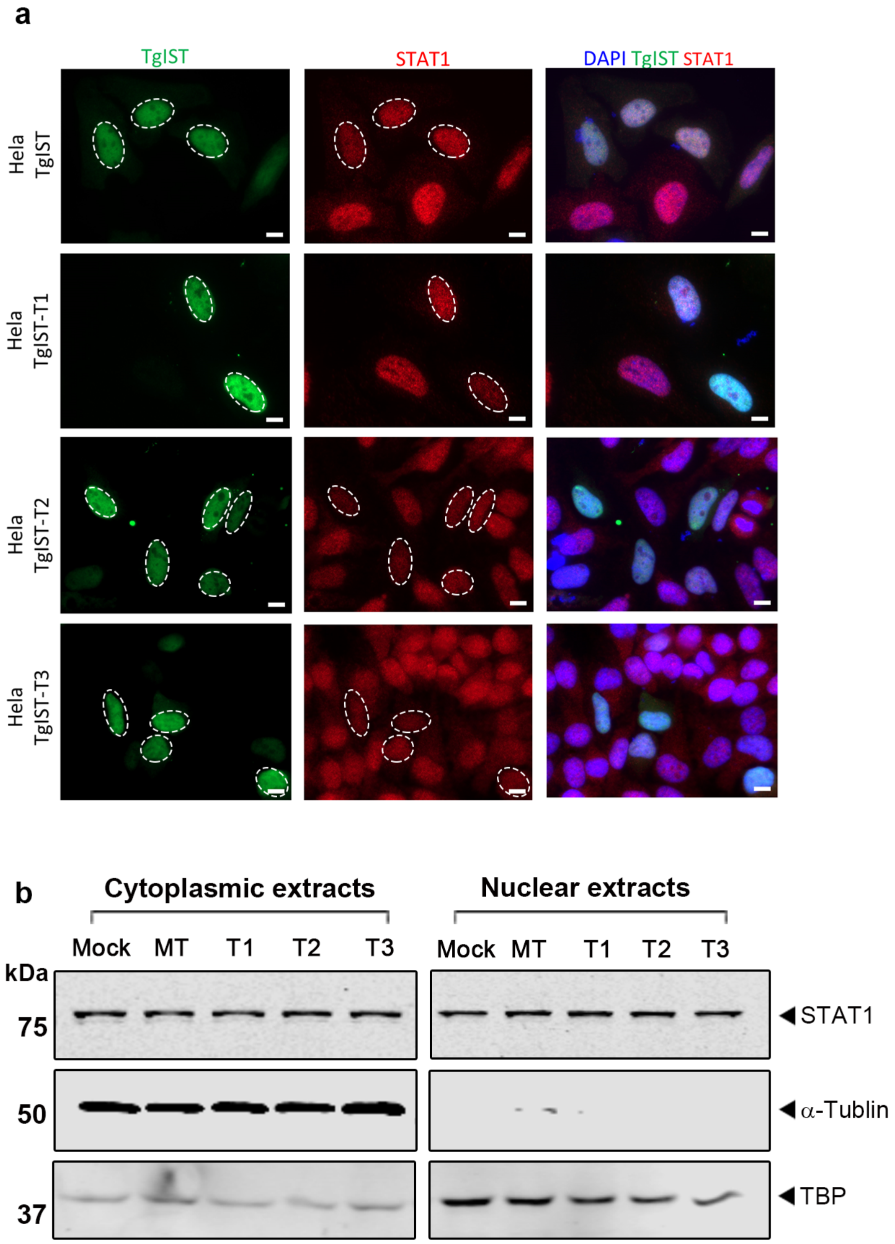


**Fig. S2**

**STAT1 levels remain unchanged in TgIST expressing HeLa cells.** (a) Representative images showing expression of STAT1 in transfected Hela cells. Hela cells transiently expressing GFP-tagged TgIST constructs for 24 hr were activated with IFN-$\gamma$ for 6 hr followed by staining for GFP (green), STAT1 (red) and DAPI (blue). Scale bar = 10 µm. (b) Western blot analysis of cells lysates from transfected HeLa cells in (A). Equal amounts of total protein were separated by SDS-PAGE, transferred to PVDF membranes, and incubated with corresponding primary antibodies as indicated. Blots for $\alpha$-Tublin and TATA-binding protein (TBP) were used as controls for the cytoplasmic and nuclear fractions, respectively.

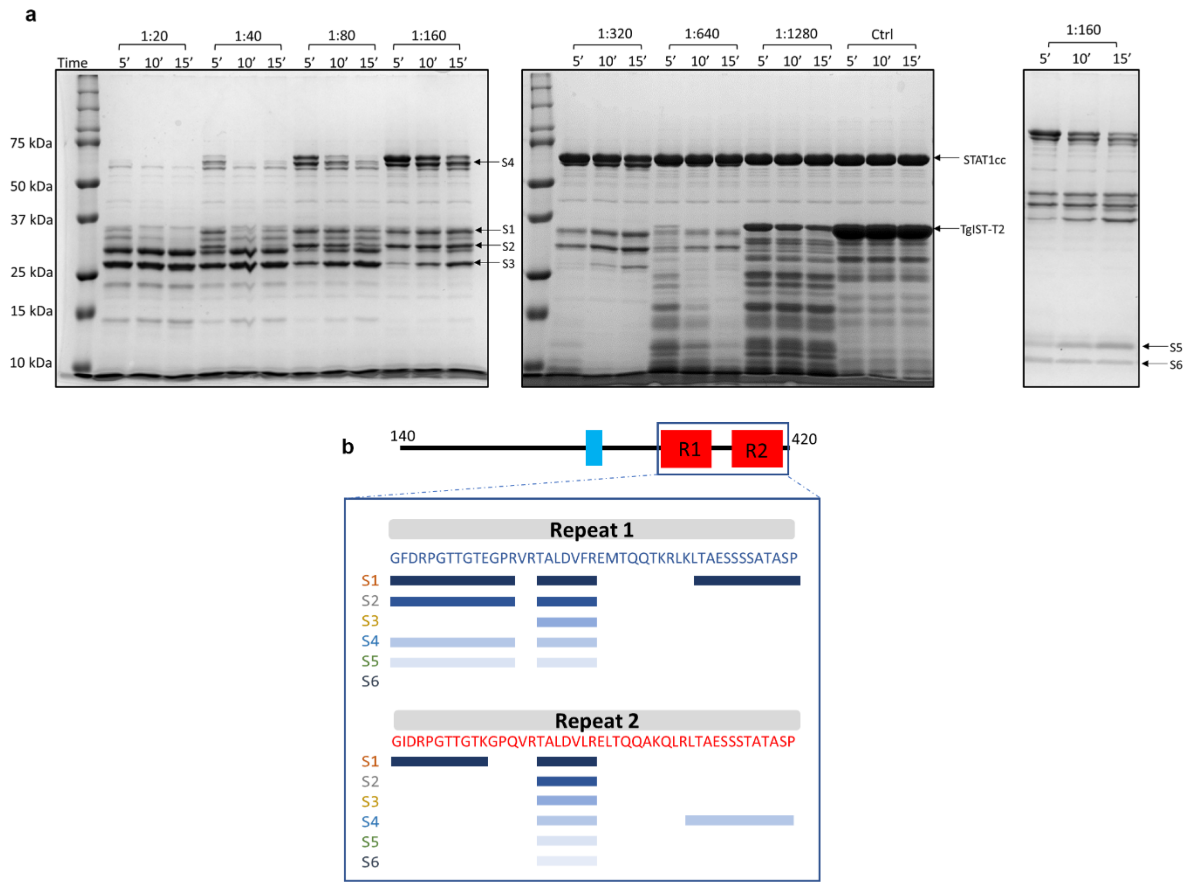


**Fig. S3**

**Identification of the core STAT1 binding sequence in TgIST by limited trypsinization and mass spectromtery. (a)** Purified TgIST-T2 complexed with STAT1cc was diluted to 10 µg in a 50 uL reaction volume. Dilutions of trypsin (1 mg/ml) from 1:20 to 1:1,280 (vol/vol %) were added to the TgIST-T2-STAT1cc complex and incubated for 5 min (5′), 10 min (10′) or 15 min (15′) as indicated. Reactions were stopped by addition of SDS sample buffer, followed by separation of samples by SDS-PAGE using 12% (the left and center gels) or 15% (the right gel) acrylamide gels. Resistant bands (numbered S1 – S6) from the samples treated with a 1:160 dilution of trypsin were cut from the gel and subjected to MS/MS analysis. (b) Limited proteolysis and mass spectrometry (MS) analysis identified core regions in the repeats of TgIST that were protected by interaction with STAT1. Purified TgIST-T2 complexed with STAT1cc was treated with trypsin and resistant bands were isolated from SDS-PAGE gels for MS analysis. Regions identified from MS are shown as rectangles below the amino acids sequence of each repeat. S1 through S6 refer to partial degradation patterns detected by SDS-PAGE.

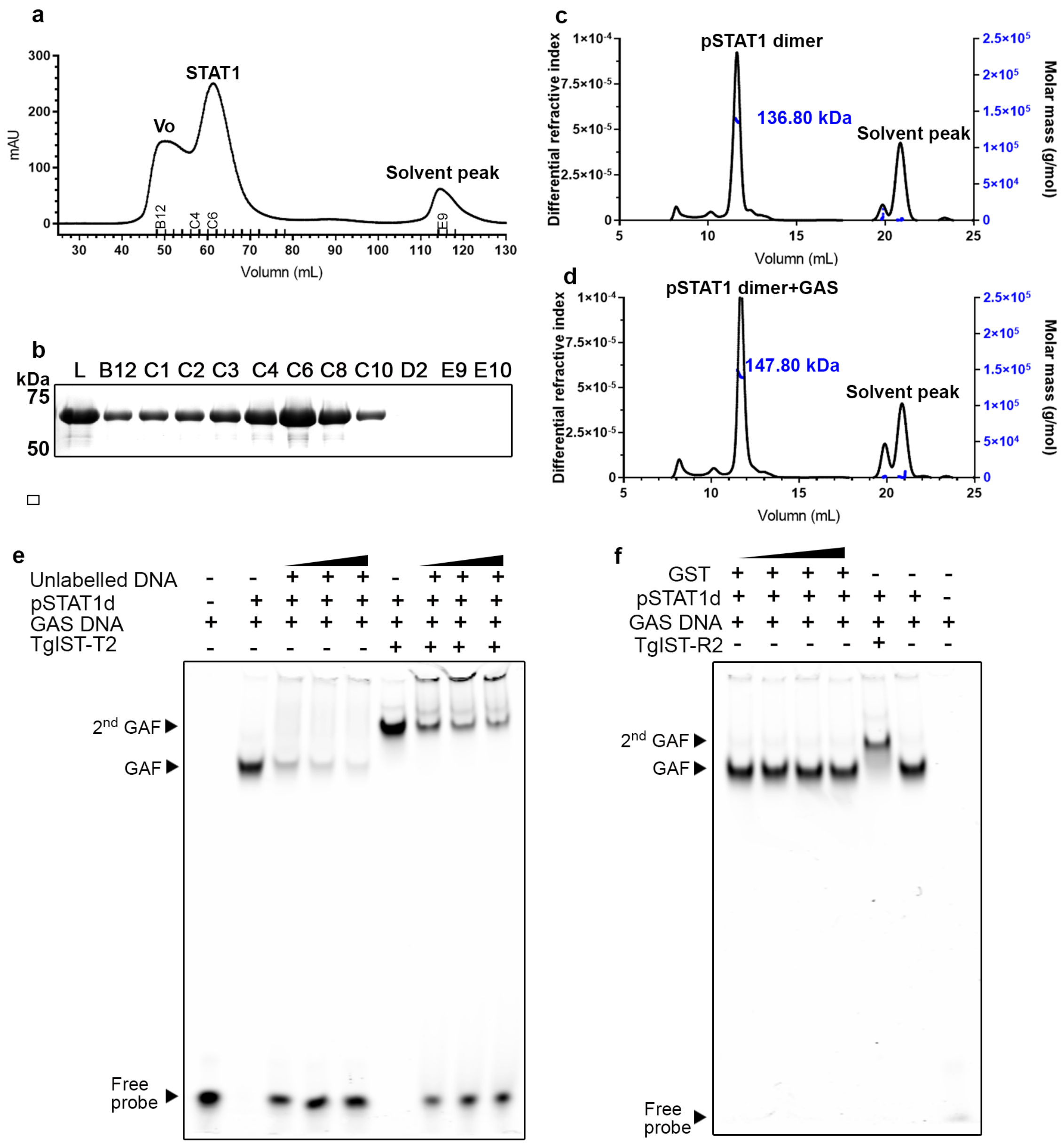


**Fig. S4**

**Purification of phosphorylated STAT1 dimer and specificity controls for EMSA assays.** (a) N-terminal Strep-tagged STAT1 was expressed in *E.coli* TKB1 cells that co-express ELK kinase in order to generate phosphorylated STAT1 dimers (pSTAT1d). pSTAT1d was successfully separated from STAT1 aggregates (Vo peak) by size exclusion chromatography. (b) SDS-PAGE analysis of column fractions from a. L, sample loaded on column. (c) Pooled fractions (C5-C10) (b) were concentrated and analyzed by multiple angle laser light scattering with in-line size exclusion chromatography (SEC-MALS).(d) Fractions C4-C10 (b) were pooled, mixed with a double stranded oligo for the GAS sequence, and subjected to SEC-MALS analysis. The observed molecular weights of pSTAT1d alone *vs*. bound to the GAS oligo were 136.8 and 147.8 kDa as measured by SEC-MALS, which compare favorably to their theoretical molecular weights of 137.0 kDa and 146.80 kDa, respectively. (e) EMSA experiments testing the specificity of formation of complexes in the absence or presence of TgIST. Addition of increasing concentrations of unlabeled DNA competitor reduced the formation of GAF in the absence of TgIST-T2 and the $2^{nd}$ GAF in the presence of TgIST-T2. GAF, gamma-activated factor, $2^{nd}$ GAF, super shifted form of GAS. (f) Addition of GST protein alone, which was used as the affinity tag in purification of TgIST-R2 in EMSA experiments shown in Figure 4, did not super shift the GAF complex. Black triangles indicate increasing concentration of components added based on label at the top.

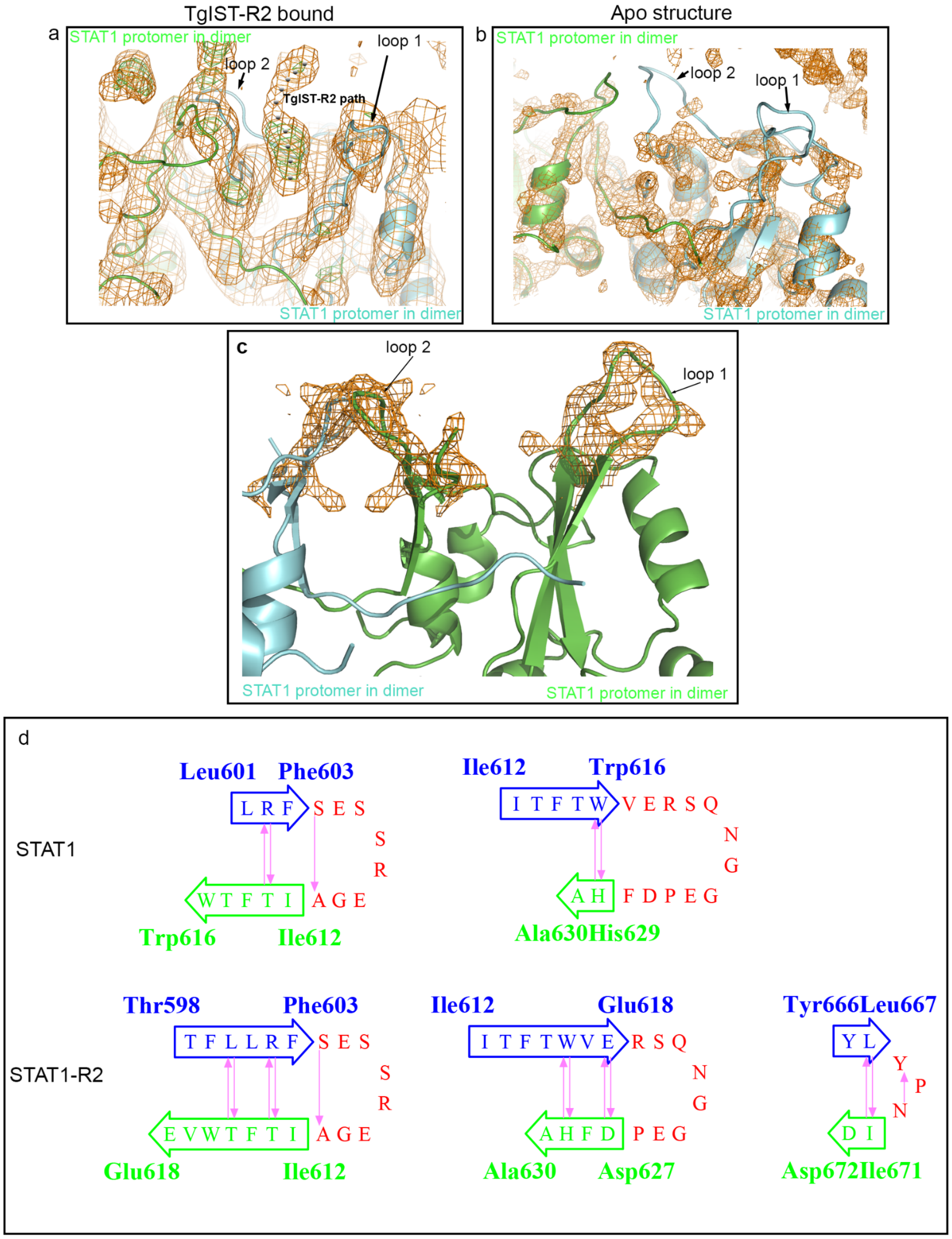


e

```
               : 1---------11--------21--------31--------41----  :
OrigSeq        : GIDRPGTTGTKGPQVRTALDVLRELTQQAKQLRLTAESSSTATASP  : OrigSeq

Jnet           : -------------HHHHHHHHHHHHHHHHHHHHHE------------  : Jnet
jhmm           : --------------HHHHHHHHHHHHHHHHHHHHE------------  : jhmm
jpssm          : -------------HHHHHHHHHHHHHHHHHHHHEEE-----------  : jpssm

Lupas 14       : --------------cccccccCCCCCCCCCCCCCC-----------  : Lupas 14
Lupas 21       : --------------CCCCCCCCCCCCCCCCCCCCCccccccc----  : Lupas 21
Lupas 28       : --------------cccccccccccccccccccccccccccc----  : Lupas 28
```

**Fig. S5**

**Electron density map of loop regions in STAT1-TgIST-R2 structure and corresponding structural analysis.** (a) Electron density map of TgIST-R2 bound to phosphorylated STAT1 dimer (pSTAT1d). The two proteins were purified separately and crystallized together using a molar ratio of 1:2.1 (pSTAT1d: TgIST-R2). An additional density is seen located at the top of the pSTAT1d interface formed by two flexible loops (loop 1 and loop 2). Orange mesh represents the 2Fo-Fc map contoured at 1 $\sigma$, Green mesh represents the Fo-Fc map contoured at 3 $\sigma$. Black dots represents the putative TgIST-R2 binding path between loop1 and loop 2.

(b) Electron density map of the structure of phosphorylated STAT1 dimer (pSTAT1d). Orange mesh as defined in (a) while no additional density is seen between the two loops. (c) Side view of additional densities located at the top of the pSTAT1d interface formed by two flexible loops. Orange mesh represents the 2Fo-Fc map contoured at 1.5 $\sigma$. (d) Wiring diagram of the $\beta$-sheet and hairpin analysis in the C-terminal STAT1 or STAT1-R2 structure. Figure was generated using PDBSum (http://www.ebi.ac.uk/thornton-srv/databases/cgi-bin/pdbsum/). Strand 1 (blue) and strand 2(green) form a hairpin shown in red. Main chain hydrogen bonds are indicated in purple. (e) Secondary structure prediction for TgIST-R2 using I-tasser server (https://zhanglab.ccmb.med.umich.edu/I-TASSER/) indicates the repeat region has a tendency to form an alpha helix. H stands for helix, C stands for coiled-coil.

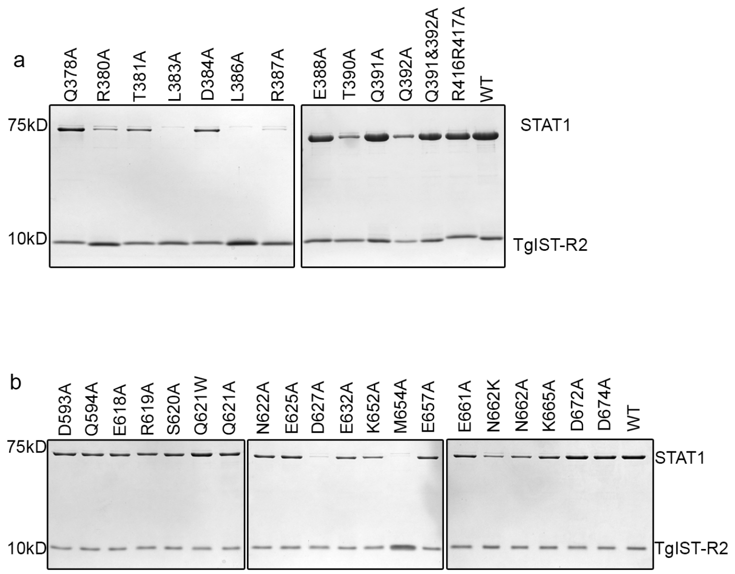


**Fig. S6**

**Binding of His-tagged TgIST-R2 to STATcc dimer assessed by nickel affinity purification.** (a) His-tagged TgIST-R2 and mutants were tested by copurification with STATcc. Mutants in TgIST are denoted above the gel. (b) His-tagged TgIST-R2 was tested by copurification with STATcc and its mutants. Mutants of STAT1cc are denoted above the gel. Eluted fractions were separated by SDS-PAGE gel and stained with Coomassie blue. One set of representative gels are shown here.

a

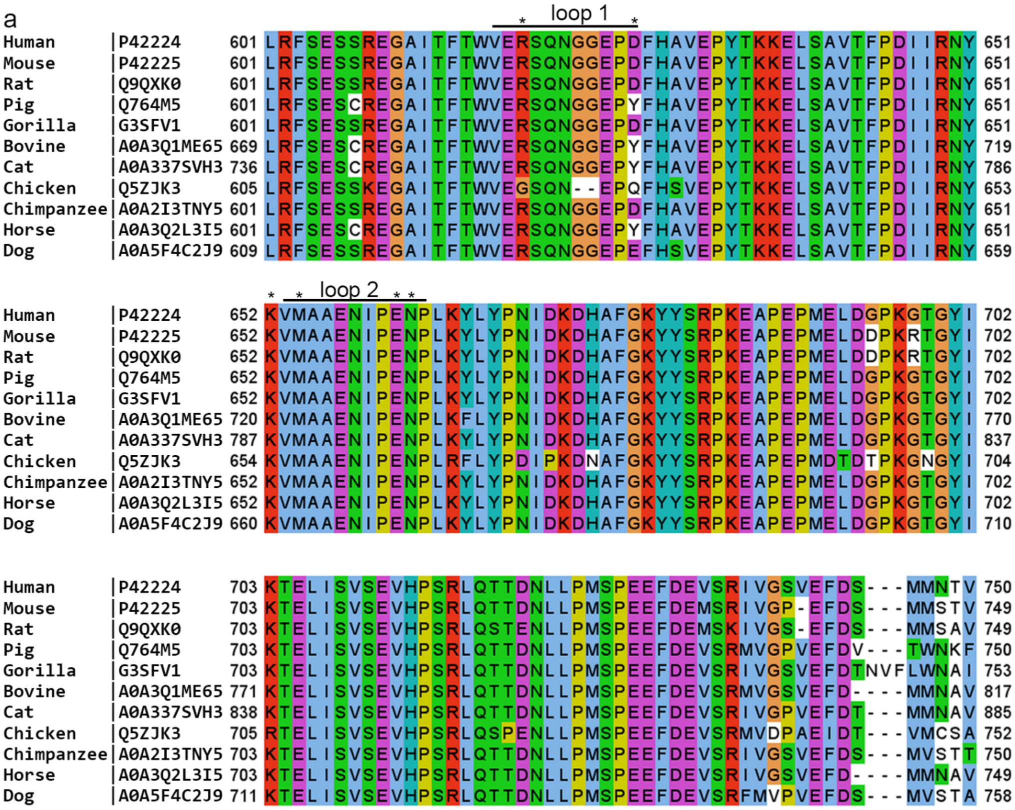


b

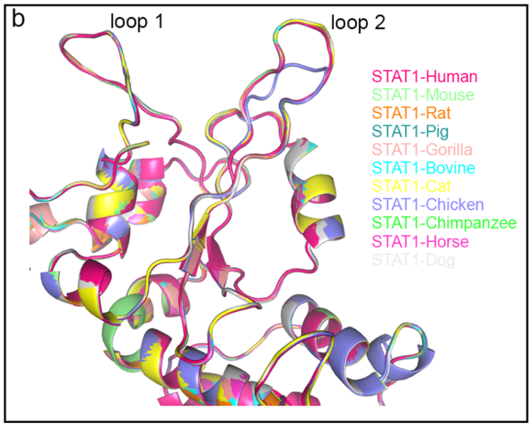


c

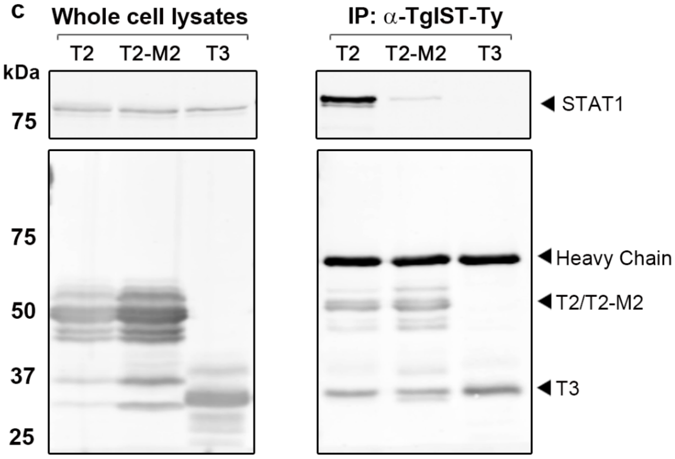


d

| | | Experiment 1 | | | Experiment 2 | | |
|---|---|---|---|---|---|---|---|
| | | HEK293T transfected with TgIST constructs: | | | | | |
| | | TgIST-T2 | TgIST-T2-M2 | TgIST-T3 | TgIST-T2 | TgIST-T2-M2 | TgIST-T3 |
| **Scaled protein abundance** | **STAT1** | 143.3 | 116 | 111.2 | 62.2 | 93.6 | 99.2 |
| | **TgIST-T2** | 374.7 | 23.8 | 1.4 | 363.3 | 35.9 | 0.7 |
| | **CBP+p300** | 82.5 | 171.6 | 105.6 | 57.8 | 147.8 | 83.8 |

e

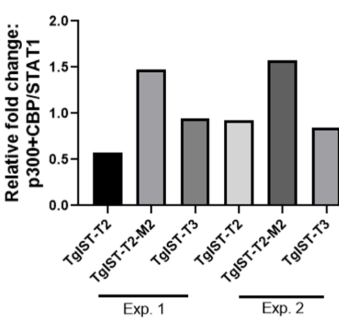

**Fig. S7**

**The SH2 domain is conserved among STAT1 from different species and western blot analysis of proteins following immunoprecipitation of TgIST-Ty from HEK293T cells.** Multiple sequence alignment of the STAT1 proteins from different species (a) and the superposition of their corresponding homology models (b). (a) STAT1 protein sequences were retrieved and imported into Jalview software (https://www.jalview.org/) and the multiple sequence alignment was performed by Muscle ( https://www.ebi.ac.uk/Tools/msa/muscle/) using default settings. The sequence accession numbers of STAT1 in Uniprot database (https://www.uniprot.org/ ) after the species name to the left of each sequence in the alignment. Color schemes are as follows, blue: hydrophobic residues; red: positively charged residues; magenta: negatively charged residues; green: polar residues; pink: cysteines; orange: glycines; yellow: prolines; and cyan: aromatic residues. (b) Homology models of the STAT1 sequences described above were built using the Swiss-model server (https://swissmodel.expasy.org/ ) based on the STAT1 dimer structure (PDB: 1BF5) as the template. Models were aligned and visualized by Pymol (https://pymol.org/2/ ). (c) HEK293T cells were transfected with TgIST constructs expressing the mature form of TgIST (M2), a truncated form containing both repeat TgIST-T2 (T2); a mutant where the core 7 amino acids in both repeats have been replaced with alanine TgIST-T2-M2 (T2-M2); and a truncated version lack both repeats TgIST-T3, also see schematic in Fig. 1d and 3b). Cells were infected for 23 hr, then treated ± IFN-$\gamma$ (100 U/mL) for additional 60 min prior to whole cell extract preparation. Membranes were incubated with corresponding primary antibodies as indicated and then IR dye-conjugated secondary antibodies. Visualization was performed using an Odyssey infrared imager. (d) Label-free quantification by mass spectrometry of STAT1 immunoprecipitation (IP) from TgIST transfected HEK293T cells, corresponding to Fig. 7e. The peptides were quantified using the precursor abundance based on intensity. Then proteins were scaled using total peptide amount. (e) Relative fold change of CBP/p300 calculated from (d). Relative fold change was defined by using scaled abundance of CBP+p300 to divide the abundance of STAT1 in each sample.
